# Characterization of metabolic compartmentalization in the liver using spatially resolved metabolomics

**DOI:** 10.1101/2021.12.13.472421

**Authors:** Jiska van der Reest, Sylwia A. Stopka, Walid M. Abdelmoula, Daniela F. Ruiz, Shakchhi Joshi, Alison E. Ringel, Marcia C. Haigis, Nathalie Y. R. Agar

## Abstract

Cells adapt their metabolism to physiological stimuli, and metabolic heterogeneity exists between cell types, within tissues, and subcellular compartments. The liver plays an essential role in maintaining whole-body metabolic homeostasis and is structurally defined by metabolic zones. These zones are well-understood on the transcriptomic level, but have not been comprehensively characterized on the metabolomic level. Mass spectrometry imaging (MSI) can be used to map hundreds of metabolites directly from a tissue section, offering an important advance to investigate metabolic heterogeneity in tissues compared to extraction-based metabolomics methods that analyze tissue metabolite profiles in bulk. We established a workflow for the preparation of tissue specimens for matrix-assisted laser desorption/ionization (MALDI) MSI and achieved broad coverage of central carbon, nucleotide, and lipid metabolism pathways. We used this approach to visualize the effect of nutrient stress and excess on liver metabolism. Our data revealed a highly organized metabolic compartmentalization in livers, which becomes disrupted under nutrient stress conditions. Fasting caused changes in glucose metabolism and increased the levels of fatty acids in the circulation. In contrast, a prolonged high-fat diet (HFD) caused lipid accumulation within liver tissues with clear zonal patterns. Fatty livers had higher levels of purine and pentose phosphate related metabolites, which generates reducing equivalents to counteract oxidative stress. This MALDI MSI approach allowed the visualization of liver metabolic compartmentalization at high resolution and can be applied more broadly to yield new insights into metabolic heterogeneity *in vivo*.

## Introduction

Advances in single-cell analysis approaches have revealed that cells within tissues can be metabolically distinct and have unique contributions to physiology and pathology (1). Metabolic compartmentalization between cellular organelles, within organs, and at the whole-body level is essential to meet the bioenergetic and anabolic demands of organisms. Additionally, metabolically distinct microenvironments develop within tissues based on physiological factors such as proximity to vasculature, which supplies nutrients and oxygen while removing metabolic waste products.

The liver is organized by regions of functional and spatial heterogeneity. Hepatocytes are structured in neat rows along the liver lobule axis from the portal vein that receives venous blood from the gut towards the central vein, which returns the blood into circulation (2). As such, the oxygen gradient is highest for periportal hepatocytes and decreases towards the pericentral area (3). Opposing gradients of oxygen and Wnt signaling along with the radial lobule axis drive differential gene expression signatures (4): approximately half of all genes in mouse hepatocytes are expressed in a zonated fashion in both space and time (5–7). This organization drives profound differences in metabolism: periportal hepatocytes rely on the oxidation of fatty acids for energy and perform metabolic functions such as gluconeogenesis, the urea cycle, and biosynthesis of cholesterol and proteins (8). In contrast, pericentral hepatocytes display glycolytic energy metabolism and synthesize lipids, bile, and glutamine.

The liver plays an essential role in maintaining whole-body metabolic homeostasis in response to nutrient abundance and restriction (9). In a satiated state, hepatocytes oxidize glucose to generate energy and synthesize fatty acids (10). Fatty acids are then esterified into triacylglycerols (TAGs) and transported to the adipose tissue for storage. In fasted conditions, the adipose tissue releases fatty acids for oxidation by the liver to yield ketone bodies that can fuel distant organs (11). Additionally, the liver performs glycogenolysis and gluconeogenesis to restore circulating glucose levels upon fasting. In contrast, upon prolonged nutrient excess conditions, the liver acts as an overflow depot for lipids when the endocrine and storage functions of the adipose tissue become compromised (12). With rising rates of obesity, nonalcoholic fatty liver disease (NAFLD) is an increasing cause of morbidity and mortality.

Despite the liver’s central role in metabolic homeostasis, liver metabolism is characterized mostly on the gene, protein, and signaling levels. However, as hepatocytes make up over 80% of liver mass (13), metabolite profiles obtained with conventional extraction-based metabolomic methods skew towards hepatocellular metabolism at the expense of other resident cell types. Spatially resolved metabolite profiling could yield new insights into metabolic heterogeneity and functional specialization within the liver.

Matrix-assisted laser desorption/ionization (MALDI) mass spectrometry imaging (MSI) is a label-free technique that allows for *in situ* spatial mapping and quantification of hundreds of metabolites from a single tissue section (14–16). Recent mass spectrometric advances have led to an increasingly higher spatial resolution that now approximates single-cell and sub-cellular analytic capability (17, 18). However, several outstanding challenges in sample preparation and data acquisition need to be addressed to ensure the robustness of metabolome-scale analyses (19, 20). Unique adaptations are required to yield reproducible and biologically relevant data for small metabolite analyses, including quenching metabolic activity, metabolite stabilization, matrix optimization, and data acquisition (14).

In this study, we implemented MALDI MSI to spatially map the distribution of small metabolites to faithfully recapitulate key bioenergetic activities. We interrogated the liver metabolic response to nutrient stress and excess conditions with a spatial resolution of identified patterns of metabolic specialization within liver tissues. We observed that fasting-induced fuel switching in the liver while in conditions of prolonged nutrient excess induced by a high-fat diet, mice develop fatty livers that remodel central carbon metabolism towards increased pentose phosphate pathway and purine metabolism. Taken together, we show that introducing spatiality into metabolomic analyses reveals an additional layer of metabolic complexity and that our workflow can be applied broadly to yield new insights into metabolic heterogeneity *in vivo*.

## Materials and Methods

### Mouse studies

C57BL/6J (000664) and BALB/cJ (000651) mice were obtained from The Jackson Laboratory. Mice were housed at 20-22°C on a 12 h light/dark cycle with ad libitum access to food (PicoLab Rodent Diet 5053) and water. All animal studies were performed in accordance with Haigis lab protocols approved by the Standing Committee on Animals, the Institutional Animal Care and Use Committee at Harvard Medical School. For heat inactivation studies, 3 mice were used (C57BL/6J, female, 7 weeks old) and kidneys, brain halves, and liver lobes from the same individual animal were subjected to the different heat inactivation treatments (overview in Supplementary Fig. 1A, E). For desiccation experiments, 2 mice were used (C57BL/6J, male, 7 weeks old). For fasting experiments, two independent cohorts of 5 mice were used per treatment group (BALB/cJ, female, 10-11 weeks old) and mice were subjected to a 16 hour overnight fast. For HFD experiments, two independent cohorts of 4 mice were used per treatment group (C57BL/6J, female). Mice were assigned at 5 weeks old to the control diet (PicoLab Rodent Diet 5053) or HFD (Research Diets, Inc. #12492) and maintained on this diet for 4.5 months. The control diet is 4.07 Gross Energy Kcal/g. The HFD is 5.21 Kcal/g. for 8-10 weeks. Comparative MALDI MSI and LC-MS analyses of tissues were always performed on the same tissue specimens.

### Tissue isolation

Mice were anesthetized with isoflurane and sacrificed by cervical dislocation. The gall bladder was removed before livers, kidneys, and brains were harvested and carefully positioned into 15 mL flat bottom specimen vials (Nalgene, Millipore Sigma), snap-frozen in liquid nitrogen, and stored at -80 °C until further processing.

### Tissue heat inactivation

Freshly resected or snap-frozen tissues were placed in sealed Maintainor® tissue cards and placed in the Stabilizor™ system (Denator AB). Sample state was specified (frozen or fresh) and the instrument determined durations of heat treatment based on sample volume for consistent and reproducible heat treatment, according to the manufacturers instructions. Next, tissues were carefully positioned into 15 mL flat bottom specimen vials (Nalgene, Millipore Sigma), snap-frozen in liquid nitrogen, and stored at -80 °C until further processing.

### MALDI tissue preparation

Frozen tissues were placed at -20 °C before sectioning in a Microm HM550 cryostat (Thermo Scientific™). Tissues were sectioned at 10 µm thickness and thaw mounted onto indium-tin-oxide (ITO)-coated slides (Bruker Daltonics) for MALDI MSI analysis with serial sections mounted onto glass slides for histological analyses. The microtome chamber and specimen holder were maintained between -15 °C and -20 °C. Slides were stored at -80 °C until further processing. For desiccation experiments, slides were subjected to desiccation in a tabletop vacuum desiccator before freezing.

### Matrix deposition

A 1,5-Diaminonaphthalene(DAN)-HCl matrix solution was used for all experiments. To generate the hydrochloride derivative of 1,5-DAN, 39.5 mg of 1,5-DAN was dissolved in 500 µL of 1 mol/L hydrochloride solution with 4 mL HPLC-grade water. The solution was sonicated for 20 minutes to dissolve 1,5-DAN, after which 4.5 mL ethanol was added to yield the matrix solution. Matrices were deposited on slides and tissues using a TM-sprayer (HTX imaging, Carrboro, NC). DAN-HCl matrix spray conditions used where: a flow rate of 0.09 mL/min, spray nozzle temperature of 75 °C, and spray nozzle velocity of 1200 mm/min. A four-pass cycle was used with 2 mm track spacing and the nitrogen gas pressure was maintained at 10 psi. For fasting experiments, ^15^N_5_-ATP (at 10 µM), ^15^N_5_-AMP (at 1 µM), and ^15^N_-_glutamate (at 100 µM) were spiked into the matrix and used as internal calibrants.

### MALDI data acquisition

A timsTOF fleX mass spectrometer (Bruker Daltonics) was used for data collection, and data was acquired using FlexImaging 5.1 software (Bruker Daltonics). The instrument was operated in negative ion mode covering the *m/z* range of 100-1350 for heat inactivation experiments and 100-1250 for desiccation experiments; a spatial resolution of 50 µm was used to define a pixel. For fasting experiments, the instrument was operated in negative ion mode covering the *m/z* range of 50-1000 *m/z*; a spatial resolution of 30 µm was used to define a pixel. Each pixel consisted of 800 laser shots, in which the laser frequency was set to 10,000 Hz. Mass calibration was performed using the dual-source ESI option with Agilent tune mix solution (Agilent Technologies) on the optimized method; for fasting experiments, the matrix was spiked with ^15^N_5_-ATP, ^15^N_5_-AMP, and ^15^N_-_glutamate which were used as internal calibrants. The heat inactivation dataset was post-calibrated using metabolites in the 133-700 *m/z* range with Data Analysis 5.3 software (Bruker Daltonics).

### MALDI data analysis

MSI data were analyzed and visualized using SCiLS Lab 2021a software (Bruker Daltonics). Imported peaks were moved to the local max using the mean spectra with a minimal interval width of 5 mDa. Peaks were then normalized to total ion current (TIC), except for desiccation experiments, as these two datasets were acquired from separate slides and runs. Ion images for metabolites of interest were generated based on peak lists containing theoretical *m/z* and ppm errors associated with the assignments were calculated. To generate segmentation maps showing regions of spectral similarity, bisecting *k*-means clustering was applied to all individual peaks in the dataset using the correlation distance metric in SCiLS Lab 2021a software. Vascular regions were defined based on the distribution of heme B, and tissue regions based on the segmentation maps, and regions of interest (ROIs) were drawn by hand. For feature annotations, statistical analyses, and quantitation; ROIs defined in SCiLS lab were exported to MetaboScape 6.0 (Bruker Daltonics). Speckle size and number were adapted for each dataset to achieve maximum pixel coverage of equal percentage for each ROI between the experimental groups. Features were annotated based on theoretical *m/z* of 114,008 metabolite entries in the Human Metabolome Database (HMDB) version 4.0 and curated based on ppm error associated with the assignment (38). To determine which metabolite intensity changes reached statistical significance, Bucket Tables were normalized for the Sum of Buckets and a two-sided student’s t-test was performed using a significance threshold of p<0.05 and a fold change >1.5. For extravascular tissue comparisons in fasting experiments, one ROI was used per biological replicate (n=5 per group). For intravascular comparisons, one ROI was used per biological replicate (n=3 per group), where the 3 replicates were selected based on which tissue cross-sections contained vascular regions of comparable size. For other comparisons, ROIs encapsulated the full tissue section.

### Dimensionality reduction and data visualization

Dimensionality reduction was used to enable interpretable visualization of the high dimensional spectra using Uniform Manifold Approximation and Projection (UMAP) (21). The UMAP learns similarities of the mass spectra in the high-dimensional space and then projects it into a lower dimensional space of two dimensions, where similar spectra are projected close to each other and dissimilar ones are projected further away. UMAP (21) was performed in an unsupervised manner and the reduced data was then colored based on the treatment (for heat inactivation experiments) or treatment, mouse ID, or metabolite of interest (for fasting experiments). The analysis was performed in R software (version 4.0.3) using the publicly available UMAP library and visualized using ggplot2 (39).

### Pathway enrichment analysis

Pathway analysis was performed using MetaboAnalyst 4.0 (40). Metabolite features identified as significantly increased after fasting in Metaboscape were exported to MetaboAnalyst using the associated HMDB ID. The enrichment method used was a hypergeometric test and the topology analysis used was relative-betweenness centrality, with the KEGG reference library (41).

### Pathway visualization

Pathways of interest were constructed in PathVisio 3.3.0 (42)and imported into MetaboScape 6.0 (Bruker Daltonics) using the “Pathway Mapping” tool to visualize the relative changes in metabolite levels.

### Metabolite colocalization analysis

To determine colocalization of DHA and ARA, metabolite intensity plots were generated in Fiji (ImageJ 1.53c) (43). Ion images for DHA and ARA were exported from SCiLS Lab and converted to 16-bit in Fiji. Windows were synchronized and freehand lines were drawn between adjacent vessels. Metabolite intensity plots were then generated along this line, with metabolite intensity (gray value) as a function of distance between vessels (in pixels). Data were exported and visualized using GraphPad Prism 8.2.1 software (GraphPad Software).

### Metabolite extraction from tissue

Frozen tissues were maintained under dry ice vapor to remain frozen until extraction, and 10-20 mg was excised with a razor blade and samples were transferred to pre-chilled Eppendorf tubes. Extraction solution consisted of a pre-chilled (−20 °C) solution of 2:2:1 HPLC-grade acetonitrile:methanol:water with 0.1 mol/L formic acid. Pre-chilled stainless-steel beads were added to Eppendorf tubes containing tissue samples, before extraction solution was added to achieve a concentration of 20 mg/mL before immediate lysis in a benchtop TissueLyser LT (Qiagen) operated at 50 Hz for 3 minutes. Next, 15% ammonium bicarbonate solution (filtered, room temperature) was added to achieve an 8% (v/v) solution and samples were lysed for another 3 minutes at 50 Hz. Samples were transferred to a benchtop shaker and vortexed at 4 °C for 15 minutes. Beads were removed and samples were centrifuged at 16,000 × g at 4 °C for 20 minutes. Clear supernatant was transferred to glass HPLC vials for immediate HPLC-MS analysis.

### HPLC-MS analysis

An iHILIC column (HILICON) was used with SII UPLC system (Thermo Fisher Scientific) coupled to a Q-Exactive HF-X orbitrap mass spectrometer (Thermo Fisher Scientific) operated with electrospray (ESI) ionization in negative ion mode at scan range *m/z* 75–1000 and a resolution of 60,000 at *m/z* 200. Buffer conditions used were: 20mM ammonium carbonate with 0.1% ammonium hydroxide in water (buffer A) and acetonitrile (buffer B). A flow rate of 0.150 mL/min was used with the following linear gradients: 0 – 20 min gradient from 80% to 20% B; 20 – 20.5 min gradient from 20% to 80% B; 20.5 – 28 min hold at 80% B; 28 – 30 min hold to waste at 80% B. Data were acquired using Xcalibur software (Thermo Fisher Scientific) and peak areas of metabolites were determined using TraceFinder 4.1 software (Thermo Fisher Scientific). Metabolites were identified by matching mass and retention time of features to commercial metabolite standards acquired previously on our instrument. Metabolite levels were normalized to tissue weight.

### Histology

Serial sections (10 µm) were fixed and stained using hematoxylin and eosin (H&E) immediately after sectioning and imaged using a bright field microscope (Zeiss Observer Z.1, Oberkochen, Germany) equipped with a Plan-APOCHROMAT lens and AxioCam MR3 camera, using a 20× or 40× magnification. High-resolution images of whole stained tissue sections were obtained using the stitching algorithm in Zeiss ZEN imaging software.

## Results

### Broad coverage of small metabolites with spatial resolution

Several experimental parameters needed to be assessed to faithfully recapitulate tissue metabolism *in situ* to visualize regions of metabolism in the liver. As major concerns are residual enzyme activity and non-enzymatic breakdown of labile metabolites, we evaluated whether enzyme inactivation through desiccation or heat inactivation treatment would stabilize tissue metabolites for MALDI MSI sample preparation. We compared procedures of storing cryosectioned tissue on slides at -80 °C and thawing them in a vacuum desiccator to minimize rehydration due to condensation (treatment_F_, Fig. 1A) with desiccation immediately after tissue sectioning before storage (treatment_DF_). To assess tissue integrity, serial sections of liver were H&E stained for histological analysis immediately after sectioning to evaluate the effects of freezing and desiccation. Minimal differences were observed for gross tissue morphology (Fig. 1B). We used ATP stability as an indicator of postmortem enzymatic activity and labile metabolite stability, as it is used by many enzymes and is liable to degradation. Using both methods, the ATP, ADP, and AMP ion images showed comparable spatial distributions and metabolite intensities (Fig. 1C). These metabolites displayed a gradient pattern in relation to their proximity to the vasculature.

**Figure 1.**
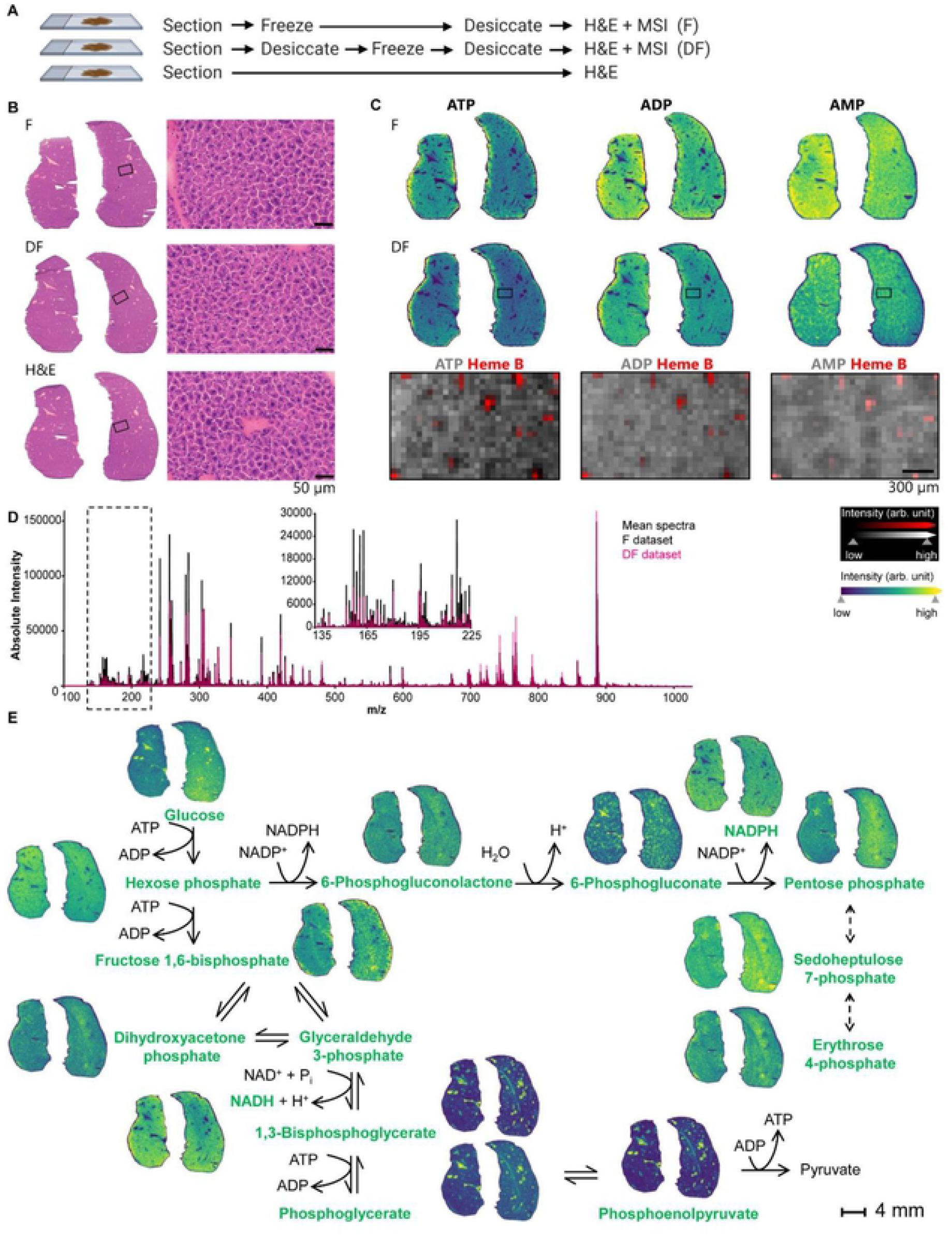
Evaluation of MALDI MSI sample preparation for small metabolites analysis. *(A)* Schematic overview of treatments where serial tissue sections were either frozen at -80 °C (treatment_F_), desiccated before freezing (treatment_DF_), or subjected to H&E staining directly after sectioning (H&E). *(B)* Histological images (20x magnification) of two mouse livers subjected to the treatments indicated in *(A). (C)* Spatial mapping (30 µm pixel) of ATP, ADP, and AMP from the two liver tissue sections that underwent the treatments indicated in *(A)*. MSI ion images showing relative distribution of ATP, ADP, and AMP individually, or in relation to the vasculature indicated by heme B. *(D)* Overlaid MALDI MSI mean spectra from serial tissue sections subjected to freezing (treatment_F_) and desiccation (treatment_DF_). Inset highlights the small metabolite range between *m/z* 135-225 for the two treatments. (*E)* Schematic overview of the connected metabolic pathways of glycolysis and the pentose phosphate pathway with corresponding ion images of the metabolites indicated in green. Glucose phosphate and fructose phosphate are indicated as hexose phosphate; phosphoglycerate and phosphoglycerate are indicated as phosphoglycerate; ribose phosphate, ribulose phosphate, and xylulose phosphate are indicated as pentose phosphate, as these are isobaric species. Dihydroxyacetone phosphate and glyceraldehyde phosphate are isobaric species as well and are visualized together.

Heat inactivation was also investigated as an alternative strategy to desiccation using kidney and brain tissues in addition to livers, as these organs have distinct anatomical features and metabolic compositions (Supplementary Figure 1A). Control mouse tissues were resected and snap-frozen in liquid nitrogen before sectioning (Freeze, treatment_F_) and compared to fresh tissues subjected to heat inactivation to denature enzymes before freezing and sectioning (Heat-Freeze, treatment_HF_). The third group of tissues was snap-frozen to preserve the metabolic state immediately upon resection and then heat-treated to denature enzymes before re-freezing and sectioning (Freeze-Heat-Freeze, treatment_FHF_). Histological analysis of kidneys showed that heat treatment disrupted tissue architecture, whereas this was preserved in control tissues (Supplementary Fig. 1B-D).

MSI was performed on serial tissue sections to evaluate the effect of heat treatment on metabolite levels. The three datasets showed a highly similar spectral coverage but pronounced differences in individual ion intensities (Supplementary Figure 2A). To visualize whether heat treatment induced changes in the metabolomes, we used Uniform Manifold Approximation and Projection (UMAP), which visualizes similarities between mass spectra projecting close together, and dissimilar spectra projected further away (21). Supplementary Figure 2B shows a clear separation of data points based on heat treatment, indicating that the applied heat treatment modified the metabolome.

The tissue distribution of ATP, ADP, and AMP after heat treatment showed a relatively stable distribution upon treatment_HF_ as compared to control tissues, but a loss of overall ATP levels upon treatment_FHF_ (Supplementary Fig. 2C). MSI ion images we used to visualize the relative spatial distribution of metabolite intensities, but these images do not inform the total metabolite pools unless the MS signal is calibrated for each metabolite. Thus, we excised representative tissue slices from the same tissues that were analyzed using MSI and determined total metabolite levels using LC-MS (Supplementary Fig. 2D). As ATP use by enzymes will lead to increased levels of AMP and ADP, successful heat stabilization of enzymes should lead to stable levels of ATP, ADP, and AMP. This comparison showed that although heat stabilization of fresh tissue seemed to maintain the spatial distribution of adenosine phosphate metabolites seen in control tissues, absolute levels of ATP decreased due to thermal destabilization.

To visualize how heat treatment affected the abundance of all detected ions in an unbiased manner, we constructed an ion segmentation map using bisecting *k*-means clustering on all identified spectra. This showed that conductive heating applied with a commercial device (Denator, Gothenburg, Sweden) set to optimze heat delivery based on frozen or fresh states and with consideration to specimen dimensions led to an overall loss of spatial localization of metabolites (Supplementary Fig. 2C and 2E). Since the whole tissues were processed, the heating profiles needed to be optimized to provide uniform heating throughout the tissue; however, this was not possible due to the tissue’s thickness. Although regional clusters of metabolites in heat-treated brains were largely maintained, they could not be accurately mapped to anatomical brain regions due to the loss of tissue morphology. Together, these results indicate that the heat treatment applied to the whole tissues prior to sectioning led to disruption of tissue structure and compromised the integrity of anatomical regions. Further optimization of the heating profile for the denaturation of enzymes and the preservation of metabolites is needed for uniform stabilization that would be compatible with spatial metabolomics workflows.

Using the treatment_DF_ followed by MALDI MSI, several additional metabolites were detected. The central carbon metabolism correlates with key energetic and biosynthetic pathways, including glycolysis and the pentose phosphate pathway (PPP) (Figure 1D). As expected, hexoses were highly abundant within the vasculature of the tissue, whereas intracellular metabolites generated from glucose were enriched in extravascular compartments rather than in the vasculature. Together, these results suggest that optimized MALDI MSI sample preparation and data acquisition workflow achieve broad coverage of small metabolites to generate reproducible spatial profiles of biologically relevant metabolic pathways.

### Distinct spatially-resolved metabolic signatures were observed in fed and fasted livers

Regions of metabolism were investigated in the liver in response to fasting by generating spatially-resolved metabolic profiles. Livers from fasted mice showed marked histological differences in hepatocyte shape due to the expected depletion of glycogen (Fig. 2A). We evaluated whether tissue metabolomes remained stable during cryosectioning, as this is a lengthy process for experiments with multiple biological replicates that need to be mounted onto the same slide for data acquisition. No significant difference was observed between total ATP, ADP, or AMP levels, indicating that these labile metabolites remained stable during cryosectioning (Fig. 2B), allowing for comparison of metabolite levels under different biological conditions. As a result, we observed that fasting led to a decrease in liver ATP content with a concomitant increase in AMP, indicative of cellular nutrient stress. Using MSI, visualizing the ion intensities distributions of adenosine phosphates showed similar results; decreasing ATP abundance and increasing AMP in hepatocellular regions within the tissue were observed (Fig. 2C). A comparison of mean spectra revealed marked differences in overall metabolite intensities between control and fasted mice (Supplementary Figure 3A). To visualize these differences in an unbiased manner, we constructed a segmentation map (Fig. 3D). This visualization showed distinct metabolic clusters within different anatomical regions of the liver and between control and fasted mice, while all biological replicates within each group clustered together (Supplementary Fig. 3C). Metabolite clusters were observed for the vasculature, hepatocytes, bile ducts, and the common bile duct. These clusters corresponded with co-registered ion images of heme B, a cofactor of hemoglobin that is enriched within the vasculature, and taurocholate, the most abundant bile acid (Fig 3D) (22, 23).

**Figure 2.**
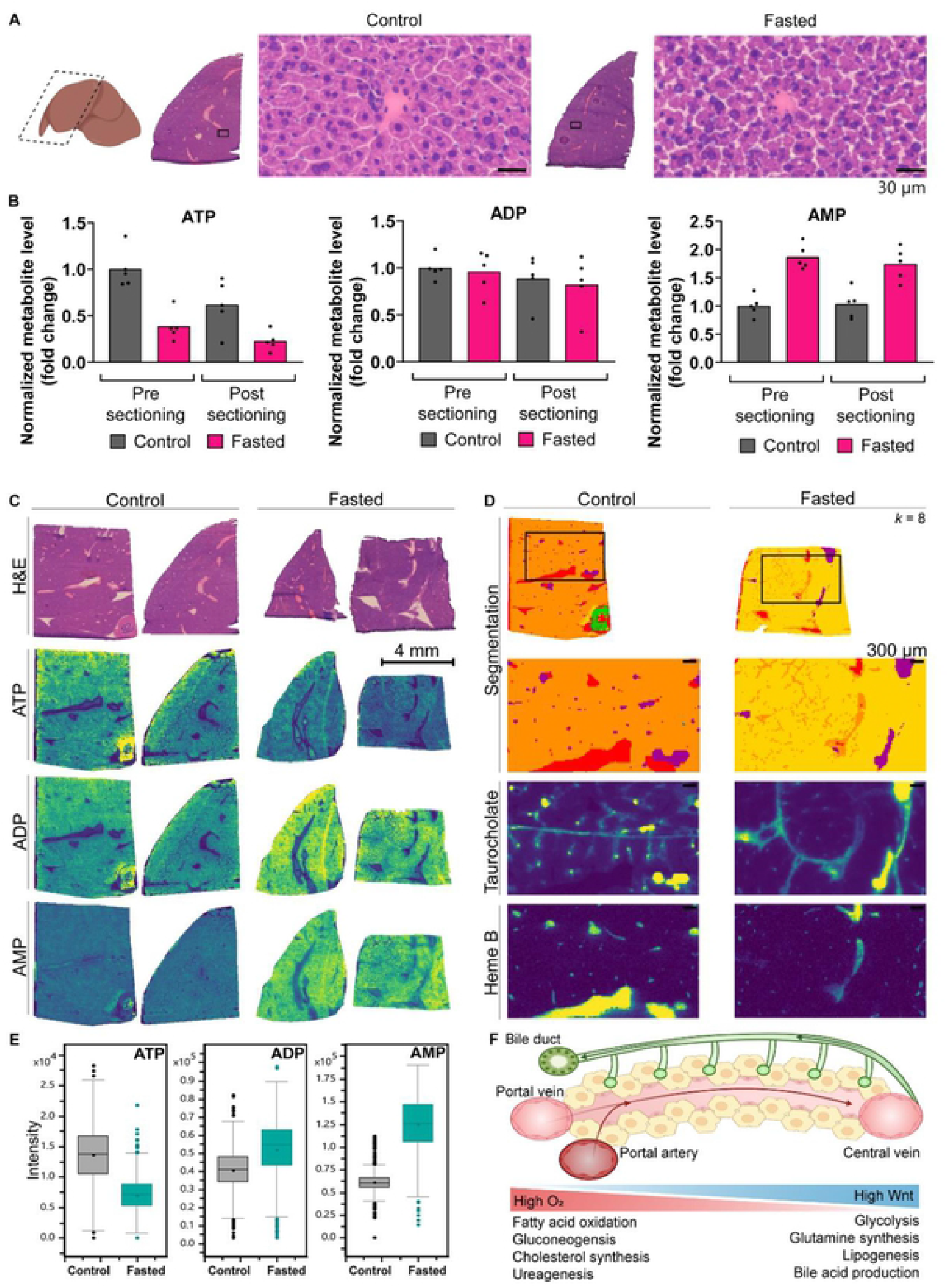
Spatially-resolved metabolic signatures in fed and fasted livers. *(A)* Histological images (40x magnification) of a representative liver section from *ad lib* fed mice and those subjected to overnight fasting; n=5 per group. (*B*) LC-MS relative quantification of total ATP, ADP, and AMP levels in liver tissues from control and fasted mice, before and after sectioning of serial sections for MALDI MSI analyses. (*C*) H&E optical and MALDI MSI ion images (30 µm pixel) of representative serial tissue sections from control and fasted mice. MSI ion images show the relative distribution of ATP, ADP, and AMP. *(D)* Segmentation map of the MALDI MSI data based on bisecting k-means clustering (*k* = 8), where each cluster is represented as an individual color, and MALDI MSI ion images of heme B as a marker of the vasculature corresponding to the red segment and taurocholate as a marker of the bile tracts corresponding to the purple segment. *(E)* MALDI MSI quantification of ATP, ADP, and AMP levels in liver tissues from control and fasted mice. *(F)* Schematic representation of metabolic gradients along the liver lobular axis. Oxygen-rich blood flows from the portal artery (dark red) and vein (light red) towards the central vein, whereas the bile (green) secreted by hepatocytes (yellow) flows through bile canaliculi in the opposite direction towards the draining bile duct. Opposing gradients of oxygen and Wnt signaling promote spatially compartmentalization metabolic functions.

**Figure 3.**
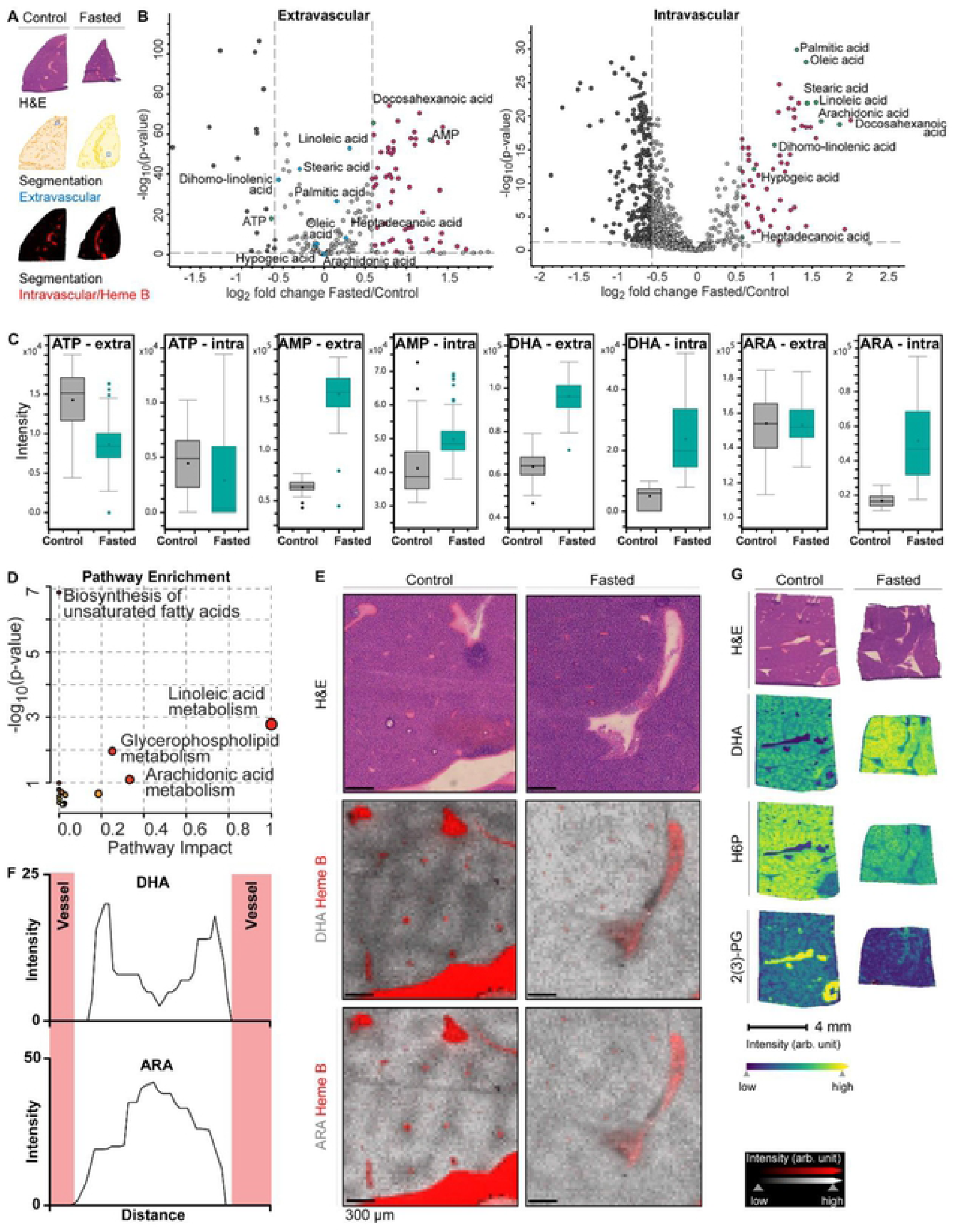
Liver metabolism and fuel switching. *(A*) H&E, segmentation, and MALDI MSI ion images of serial tissue sections from control and fasted mice indicating how extravascular and intravascular tissue regions were defined for spatial metabolic analyses. Hepatocyte-enriched regions (denoted as extravascular) were identified using the segmentation map of the MALDI MSI data based on bisecting k-means clustering (*k* = 8), with control and fasted tissues represented as dark and light orange, respectively. Intravascular regions were defined based on intensity of heme B. ROIs depicted in blue indicate where metabolite spectra were extracted for further spatial analysis. *(B)* Volcano scatterplot displaying log 2 metabolite intensity ratios vs. significance value in fasted compared to control mouse liver extravascular (left) and intravascular tissue (right). Every circle represents a unique metabolite; dark grey circles indicate metabolites depleted after fasting and magenta circles indicate metabolites enriched after fasting, that showed a fold change >1.5 between treatments and reached statistical significance (p-value <0.05). Highlighted green circles are statistically significantly changed metabolites indicating cellular energy status (AMP/ATP) and fatty acids with their corresponding names. Corresponding metabolites that were not statistically significantly changed are highlighted in blue. *(C)* MALDI MSI relative quantification of selected metabolites in the extravascular versus intravascular tissue regions. *(D)* Pathway enrichment scatterplot displaying pathway impact scores vs. significance value in fasted compared to control mouse vasculature. Increased circle size indicates pathway coverage of the identified metabolites in the dataset. Pathways identified as enriched are displayed by name. *(E)* H&E and MALDI MSI ion images (30 µm pixel) of serial tissue sections from representative control and fasted mouse livers. MSI ion images show relative distribution of DHA and ARA in relation to heme B in red, with indicated intensity scale. *(F)* Quantification of metabolite spatial distribution for DHA and ARA from blood vessel to adjacent blood vessel, where the metabolite intensity is shown as a function of distance between two vessels. Vasculature position is indicated in red. *(G)* H&E and MALDI MSI ion images (30 µm pixel) of tissue serial sections from a representative control and fasted mouse liver. MSI ion images show relative distribution of DHA, H6P, and 2(3)-PG, with indicated intensity scale.

Furthermore, using spiked internal standards into the MALDI matrix, we observed that independent mouse cohorts and replicates within each treatment group were highly reproducible in this study (Supplementary Fig. 3A-C). Together, these results indicate that our workflow yielded consistent and highly reproducible results to visualize metabolic compartments in the liver.

### Visualization of fasted liver metabolism shows disruption of metabolism and fuel switching

The liver acts as a metabolic rheostat to maintain whole-body energy homeostasis in times of nutrient stress and excess. As MSI adds a spatial dimension to metabolomic analyses, we dissected the metabolic compartmentalization in the fasting liver. We identified metabolic differences using the unbiased UMAP approach, which showed separation of data clusters from fasted compared to control livers (Supplementary Fig. 4A). Additional clusters were observed within treatment groups, visualized by coloring UMAP distributions per individual mouse (Supplementary Fig. 4B). The distribution of heme over the UMAP graphs indicated that these clusters could represent distinct anatomical regions within tissues (Supplementary Fig. 4C). To explore differences between liver and systemic metabolism, we extracted metabolite spectra from MSI data on a pixel-by-pixel basis. As shown in Figure 3A, we used the segmentation map (Fig. 2D, Supplementary Fig. 5A) to select regions-of-interest enriched for hepatocytes (extravascular tissue) or heme B (intravascular tissue, circulating metabolites). In accordance with our previous observations, ATP was significantly decreased and AMP significantly increased in extravascular tissue upon fasting (Fig. 3B, left).

We also observed an increase in the fatty acid docosahexaenoic acid (DHA). This was recapitulated in the UMAP distributions, where AMP and DHA were more abundant in fasted mice (Supplementary Fig. 4D). The metabolite profiles from intravascular regions did not show differences in adenosine phosphate metabolites, but several fatty acids were significantly enriched in the circulation upon fasting (Fig. 3B, right) whereas they were not significantly changed within extravascular regions of the tissue (Fig. C, Supplementary Figure 5C). It is well-understood that the adipose tissue releases fatty acids for oxidation by the liver to yield ketone bodies that can fuel distant organs, which is corroborated by these results and indicates that spatially-resolved metabolomics can inform on metabolic compartmentalization within tissues.

Indeed, pathway analysis of the intravascular regions showed that several lipid metabolic pathways were enriched (Fig. 3D). Interestingly, comparing the spatial distribution of fatty acids showed that the abundance of DHA and ARA follow a specific and compartmentalization pattern in fed livers (Fig. 3E, Supplementary Fig. 5B). DHA is a 22-carbon polyunsaturated omega-3 fatty acid (22:6), whereas arachidonic acid (ARA) is a 20-carbon polyunsaturated omega-6 fatty acid (20:4). Both can be synthesized from alpha-linolenic acid, which in turn is produced from the essential fatty acid linoleic acid. These fatty acids can also be released from complex lipids through lipolysis. Relative quantification of the metabolite intensity as a function of the distance between blood vessels confirmed that DHA is enriched in proximity to the vasculature while ARA displayed the opposite enrichment pattern (Fig. 3F, Supplementary Fig. 5D). Upon fasting, this distinct spatial compartmentalization within the extravascular regions is lost. In contrast to the increase in DHA within liver cells, the levels of glycolytic intermediates decreased within liver tissue, indicating a fuel switch upon fasting that decreases liver glucose use in favor of lipid metabolism. Together, these results indicate that spatially dissecting metabolite profiles can yield new insights into metabolic compartmentalization within tissues and between the local tissue environment and the circulation.

### Fatty livers show a metabolic signature indicative of oxidative stress in response to prolonged nutrient excess

We also investigated how the liver’s response to nutrient stress might contrast to its response to nutrient excess by subjecting mice to a high-fat diet (HFD). Livers of HFD mice showed marked histological differences, with hypertrophy and accumulation of lipid droplets that displayed in unique patterns where lipid droplets were deposited away from the vasculature (Fig. 4A). In human, it has been established that macrovesicular steatosis, where hepatocytes become displaced by lipid droplets, is associated with advanced fatty liver disease, inflammation, fibrosis, and poor clinical outcomes (24, 25). We evaluated the changes in metabolite levels, and subsequent pathway analysis showed that several metabolic pathways were significantly enriched upon HFD feeding, including the pentose phosphate pathway and purine metabolism (Fig. 4B). Cells increase PPP activity in response to oxidative stress to generate NADPH, a reducing factor that is essential to maintain reduced pools of glutathione, the main antioxidant in cells, and antioxidant enzymes that help maintain cellular redox balance (Fig. 4C). That HFD livers experience increased redox stress is corroborated by the observed increase in glutathione (GSH; Fig. 4C, D, E). Interestingly, although the PPP intermediates pentose 5-phosphate (P5P) and sedoheptulose 7-phosphate (S7P) are increased in fatty livers, levels of NADPH are decreased (Fig. 4D, E). This finding suggests that despite the cellular reprogramming towards an antioxidant response that occurs in fatty livers, cells have lower NADPH levels.

**Figure 4.**
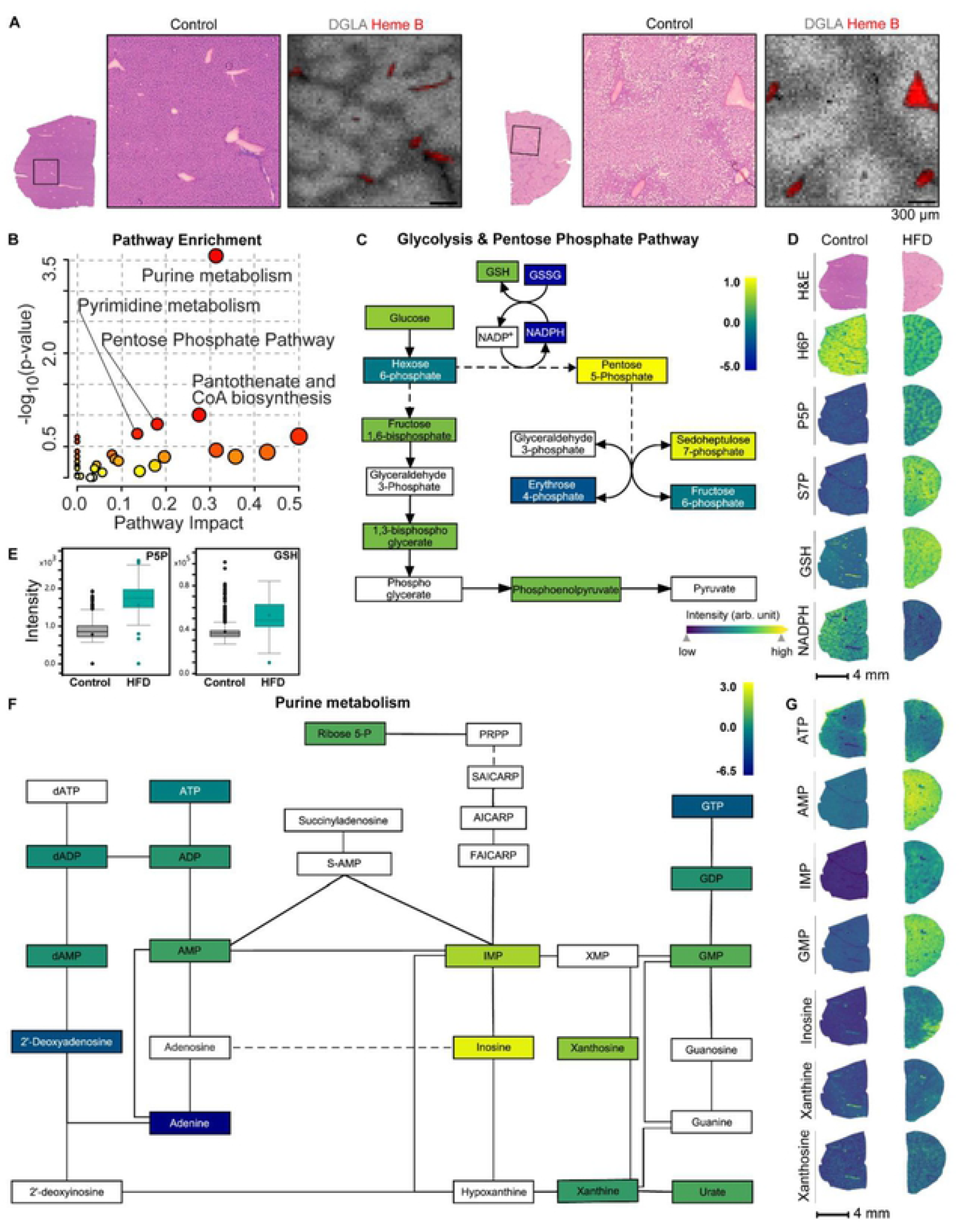
Fatty livers face oxidative stress and increased purine metabolism in response to prolonged nutrient excess. *(A*) Histological images of a representative liver section from *ad lib* fed mice on a control or high-fat diet for 4.5 months (n=5 per group, 2 independent experiments) with the corresponding ion image of the fatty acid dihomo-linolenic acid (DGLA). *(B)* Pathway enrichment scatterplot displaying pathway impact scores vs. significance value in HFD compared to control mouse liver tissues. Increased circle size indicates pathway coverage of the identified metabolites in the dataset. Pathways identified as enriched are displayed by name. *(C)* Schematic overview of the connected metabolic pathways of glycolysis and the pentose phosphate pathway with corresponding relative fold change intensities of HFD compared to control mice, with indicated intensity scale. *(D)* H&E and MALDI MSI ion images of serial tissue sections from a representative control and HFD mouse liver. The rectangles on the H&E image indicate the position of the images displayed in *(A)*. MSI ion images(30 µm pixel) show relative distribution of the indicated metabolites, with indicated intensity scale. *(E)* Absolute quantification of the indicated metabolites for control (grey) compared to HFD (green) mice. *(F)* Schematic overview of purine metabolism with corresponding relative fold change intensities of HFD compared to control mice, with indicated intensity scale. *(G)* MALDI MSI ion images of serial tissue sections from a representative control and HFD mouse liver showing relative distribution of the indicated metabolites, with indicated intensity scale.

Pathway enrichment analysis showed that in addition to the PPP, purine metabolism was significantly enriched in fatty livers (Fig. 4B). Purines are essential for supplying the building blocks for nucleotides, thereby DNA/RNA synthesis, and nucleotide cofactors such as NAD and the major energy carriers in cells (Fig. 4F, G). Increases in redox stress are known to increase DNA damage and might trigger purine metabolism to aid DNA repair, whereas the disruption of cellular energy status may converge upon the purine and pyrimidine pathways due to their important roles as cellular energy carriers to maintain cellular homeostasis. Together, these results suggest that spatially dissecting metabolite profiles and multiplexing tissue anatomical information with metabolic characterization can promote our understanding of metabolic compartmentalization in physiology and pathology.

## Discussion

Metabolic heterogeneity within tissues and metabolic crosstalk between cells are essential contributors to functional specialization in multicellular organisms. This emphasizes the need to introduce spatiality into metabolomic analyses to better understand the role of metabolic heterogeneity in physiology and disease. MALDI MSI has been used to study protein, drug and metabolite distribution in tissues from model organisms and humans to yield new biological insights. Spatially mapping endogenous metabolites can be applied to delineate metabolic properties of distinct anatomical structures (26), inform on their biological functions (27), identify abnormal or pathological regions within tissues (28), and their metabolic properties (29), and aid in surgical decision-making (30). Advances in instrumentation and application have produced increased molecular complexity and spatial resolution analyses leading to new insights into metabolic function and heterogeneity at the single-cell scale (31, 32). With increasing sensitivity and specificity in ion detection and annotation, MSI is now emerging as a tool for spatially-resolved, metabolome-scale analyses that advance our understanding of cellular and organismal biology (32). Maintaining metabolic fidelity of the tissue during sample processing is essential to yielding meaningful analyses, especially in comparison with chromatography-based mass spectrometry approaches where metabolomes are stabilized by quenching steps and samples are maintained at low temperatures until analysis while several sample preparation steps for MALDI MSI occur at ambient conditions. Here, we demonstrate an approach to prepare tissue samples for MSI that minimizes conversion or breakdown of labile metabolites while broadening the range of small metabolites detected to more broadly cover metabolic pathways and yield new insights into tissue metabolism.

Liver zonation is well-understood on the transcript level (3, 5, 6, 9, 33–35), but has not been comprehensively visualized on the metabolite level. An important advance of profiling metabolic heterogeneity on the metabolite rather than transcript level is that an immediate snapshot of metabolism can be captured instead of indirect measures provided by enzyme transcripts or protein levels. Direct metabolite profiling is enabled by the fact that MALDI MSI requires minimal sample handling, and dissociation of distinct cell types is not necessary. We were able to validate and visualize metabolic compartmentalization in liver tissues in distinct nutrient stress and excess conditions. We observed distinct metabolic profiles within zones and between tissue compartments, which may be obscured in extraction-based metabolomic analyses as the hepatocyte fraction contributes most of the mass and metabolic content of the liver. By analyzing metabolite spectra from distinct extra- and intravascular regions, we observed specific metabolic profiles consistent with the known metabolic function of each compartment. We observed a strong enrichment of fatty acids in blood vessels, consistent with the liver’s function of converting fatty acids released from the adipose tissue to generate alternative fuels for distant organs. In addition to compartmentalization between the liver organ environment and the circulation, we also observed distinct patterns of metabolite abundance within the tissue microenvironment, with hepatocytes showing enrichment of specific fatty acids based on their proximity to the vasculature. This distinct pattern was highly organized and reproducible between biological replicates in nutrient-replete conditions but vanished when facing nutrient stress after fasting. This suggests that prolonged nutrient stress induces metabolic adaptations that overrule the functional compartmentalization of hepatocytes seen under nutrient-replete conditions. In prolonged nutrient excess conditions induced by a high-fat diet, lipid droplets accumulate in the liver, forming distinct lipid depots throughout the tissue.

In contrast to fasting conditions, where glycolytic metabolism were low, fatty livers displayed higher levels of glycolytic and PPP metabolites. Together with the marked increase in GSH levels, indicates levels of oxidative stress, which is constant with high levels of NADPH levels. Additionally, we observed an increase in purine metabolism, which may produce nucleotides needed to repair DNA damage, generate essential energy carriers, or provide precursors for metabolic cofactors such as NAD, which can all become disturbed by cellular redox stress. These results indicate that although the lipid content of the liver increases upon HFD feeding, the lipid droplets act as an overflow depot rather than being effectively metabolized by the liver to dissipate excess energy. Adding a temporal component to our spatial metabolomic analyses and multiplexing with orthogonal modes of single-cell tissue imaging analyses (36, 37) may help further elucidate which regulatory nodes govern the observed fuel switching in fasting and fatty liver. Taken together, our described workflow enables the detection of endogenous metabolites and achieves a broad coverage of the tissue metabolome that can be applied to characterize and interrogate metabolic heterogeneity in physiology and pathology.

## Conclusions

Cellular metabolism is spatiotemporally heterogeneous, yet leading metabolomics approaches do not preserve spatial information. We present a MALDI MSI approach to map metabolic heterogeneity in the liver in nutrient replete, stress, and excess conditions. Our data validate and extend what is known about liver metabolic compartmentalization and visualize this at high resolution with broad coverage of key pathways in central energy metabolism. The label-free molecular imaging approach demonstrated here can be applied broadly to study metabolism in tissues and reveal new insights into metabolic heterogeneity *in vivo* to better understand the role of metabolism in physiology and pathology.

## Acknowledgments

The authors thank members of the Haigis and Agar labs for helpful discussions, in particular Dr. Elisa York for assistance with matrix calibration and Dr. Ilaria Elia and Giulia Notarangelo for helpful comments on the manuscript. J.vd.R is supported by a Postdoctoral Fellowship from the Human Frontier Science Program (LT000530/2020-L). S.A.S. is supported by an NIH T32 (award number: T32EB025823) fellowship. D.R. is supported by the NCI CaNCURE grant (award number: R25 CA174650). A.E.R is supported by postdoctoral fellowships from the American Cancer Society (United States; 130373-PF-17-132-01-CCG) and Cell Biology Education and Fellowship Fund (United States; Harvard Medical School). M.C.H. is supported by the Ludwig Center at Harvard Medical School, the Paul F. Glenn Foundation for Medical Research, and NIH grant R01DK127278. This work was funded in part by the Pediatric Low-Grade Astrocytoma Program (award #9616692) at PBTF (N.Y.R.A.). The work was also funded by NIH U54 CA210180 MIT/Mayo Physical Science Oncology Center for Drug Distribution and Drug Efficacy in Brain Tumors (N.Y.R.A.) and the Ferenc Jolesz Advanced Technologies National Center for Image Guided Therapy NIH P41 EB028741. Schematic figures were created with BioRender.com.

## Notes

**Conflict of Interest Statement** In compliance with Harvard Medical School and Partners Healthcare guidelines on potential conflict of interest, we disclose that N.Y.R.A. is scientific advisor to BayesianDx and inviCRO, and key opinion leader to Bruker Daltonics; M.C.H. has received funding from Roche.

### Competing Interest Statement

In compliance with Harvard Medical School and Partners Healthcare guidelines on potential conflict of interest, we disclose that N.Y.R.A. is scientific advisor to BayesianDx and inviCRO, and key opinion leader to Bruker Daltonics M.C.H. has received funding from Roche.

## References

1. E. K. Neumann, T. D. Do, T. J. Comi, J. v. Sweedler, Exploring the Fundamental Structures of Life: Non-Targeted, Chemical Analysis of Single Cells and Subcellular Structures. Angewandte Chemie - International Edition 58, 9348–9364 (2019).

2. S. Ben-Moshe, S. Itzkovitz, Spatial heterogeneity in the mammalian liver. Nature Reviews Gastroenterology and Hepatology 16, 395–410 (2019).

3. C. P. S. C. C Torre, Molecular determinants of liver zonation. Prog. Mol. Biol. Transl Sci. 97, 127–150 (2010).

4. C. Torre, C. Perret, S. Colnot, Transcription dynamics in a physiological process: β-Catenin signaling directs liver metabolic zonation. International Journal of Biochemistry and Cell Biology 43, 271–278 (2011).

5. K. B. Halpern, Single-cell spatial reconstruction reveals global division of labour in the mammalian liver. Nature 542, 352–356 (2017).

6. C. Droin, et al., Space-time logic of liver gene expression at sub-lobular scale. Nature Metabolism 3, 43–58 (2021).

7. S. Annunziato, J. S. Tchorz, Liver zonation—a journey through space and time. Nature Metabolism 3, 7–8 (2021).

8. N. K. K. J. D Sasse, Functional heterogeneity of rat liver parenchyma and of isolated hepatocytes. FEBS Lett. 57, 83–88 (1975).

9. L. Rui, Energy metabolism in the liver. Comprehensive Physiology 4, 177–197 (2014).

10. L. O. Schwen, et al., Zonated quantification of steatosis in an entire mouse liver. Computers in Biology and Medicine 73, 108–118 (2016).

11. M. Defour, G. J. E. J. Hooiveld, M. van Weeghel, S. Kersten, Probing metabolic memory in the hepatic response to fasting. Physiological Genomics 52, 602–617 (2020).

12. K. Qureshi, G. A. Abrams, Metabolic liver disease of obesity and role of adipose tissue in the pathogenesis of nonalcoholic fatty liver disease. World Journal of Gastroenterology 13, 3540–3553 (2007).

13. P. Godoy, Recent advances in 2D and 3D in vitro systems using primary hepatocytes, alternative hepatocyte sources and non-parenchymal liver cells and their use in investigating mechanisms of hepatotoxicity, cell signaling and ADME. Arch. Toxicol. 87, 1315–1530 (2013).

14. A. R. Buchberger, K. DeLaney, J. Johnson, L. Li, Mass Spectrometry Imaging: A Review of Emerging Advancements and Future Insights. Analytical Chemistry 90, 240–265 (2018).

15. K. A. Zemski Berry, et al., MALDI imaging of lipid biochemistry in tissues by mass spectrometry. Chemical Reviews 111, 6491–6512 (2011).

16. R. J. A. Goodwin, Z. Takats, J. Bunch, A Critical and Concise Review of Mass Spectrometry Applied to Imaging in Drug Discovery. SLAS Discovery 25, 963–976 (2020).

17. J. M. Spraggins, et al., High-Performance Molecular Imaging with MALDI Trapped Ion-Mobility Time-of-Flight (timsTOF) Mass Spectrometry. Analytical Chemistry 91, 14552–14560 (2019).

18. L. A. McDonnell, et al., Subcellular imaging mass spectrometry of brain tissue. Journal of Mass Spectrometry 40, 160–168 (2005).

19. T. J. A. Dekker, et al., Towards imaging metabolic pathways in tissues. Analytical and bioanalytical chemistry 407, 2167–2176 (2015).

20. T. M. J. Evers, et al., Deciphering Metabolic Heterogeneity by Single-Cell Analysis. Analytical Chemistry 91, 13314–13323 (2019).

21. L. McInnes, J. Healy, J. Melville, UMAP: Uniform Manifold Approximation and Projection for Dimension Reduction. arXiv (2018) (December 7, 2020).

22. H. R. Brown, et al., Drug-induced Liver Fibrosis: Testing Nevirapine in a Viral-like Liver Setting Using Histopathology, MALDI IMS, and Gene Expression. Toxicologic Pathology 44, 112–131 (2016).

23. I. Rzagalinski, N. Hainz, C. Meier, T. Tschernig, D. A. Volmer, MALDI Mass Spectral Imaging of Bile Acids Observed as Deprotonated Molecules and Proton-Bound Dimers from Mouse Liver Sections. Journal of the American Society for Mass Spectrometry 29, 711–722 (2018).

24. N. Chalasani, et al., Relationship of steatosis grade and zonal location to histological features of steatohepatitis in adult patients with non-alcoholic fatty liver disease. Journal of Hepatology 48, 829–834 (2008).

25. H. Alamri, et al., Mapping the triglyceride distribution in NAFLD human liver by MALDI imaging mass spectrometry reveals molecular differences in micro and macro steatosis. Analytical and Bioanalytical Chemistry 411, 885–894 (2019).

26. D. Trede, et al., Exploring three-dimensional matrix-assisted laser desorption/ionization imaging mass spectrometry data: Three-dimensional spatial segmentation of mouse kidney. Analytical Chemistry 84, 6079–6087 (2012).

27. E. K. Neumann, et al., Spatial Metabolomics of the Human Kidney using MALDI Trapped Ion Mobility Imaging Mass Spectrometry. Analytical Chemistry (2020) https://doi.org/10.1021/acs.analchem.0c02051 (October 13, 2020).

28. J. Wang, et al., MALDI-TOF MS imaging of metabolites with a N -(1-Naphthyl) ethylenediamine dihydrochloride matrix and its application to colorectal cancer liver metastasis. Analytical Chemistry 87, 422–430 (2015).

29. E. C. Randall, et al., Localized metabolomic gradients in patient-derived xenograft models of glioblastoma. Cancer Research 80, 1258–1267 (2020).

30. S. S. Basu, et al., Rapid MALDI mass spectrometry imaging for surgical pathology. npj Precision Oncology 3 (2019).

31. J. M. Spraggins, et al., High-Performance Molecular Imaging with MALDI Trapped Ion-Mobility Time-of-Flight (timsTOF) Mass Spectrometry. Analytical Chemistry 91, 14552–14560 (2019).

32. I. S. Gilmore, S. Heiles, C. L. Pieterse, Metabolic Imaging at the Single-Cell Scale: Recent Advances in Mass Spectrometry Imaging. Annual Review of Analytical Chemistry 12, 201–224 (2019).

33. S. Ben-Moshe, S. Itzkovitz, Spatial heterogeneity in the mammalian liver. Nature Reviews Gastroenterology and Hepatology 16, 395–410 (2019).

34. R. Gebhardt, M. Matz-Soja, Liver zonation: Novel aspects of its regulation and its impact on homeostasis. World journal of gastroenterology 20, 8491–504 (2014).

35. T. K. K Jungermann, Role of oxygen in the zonation of carbohydrate metabolism and gene expression in liver. Kidney Int. 51, 402–412 (1997).

36. E. K. Neumann, K. v. Djambazova, R. M. Caprioli, J. M. Spraggins, Multimodal Imaging Mass Spectrometry: Next Generation Molecular Mapping in Biology and Medicine. Journal of the American Society for Mass Spectrometry (2020) https://doi.org/10.1021/jasms.0c00232 (December 15, 2020).

37. J. R. Lin, et al., Highly multiplexed immunofluorescence imaging of human tissues and tumors using t-CyCIF and conventional optical microscopes. eLife 7 (2018).

38. D. S. Wishart, et al., HMDB 4.0: The human metabolome database for 2018. Nucleic Acids Research 46, D608–D617 (2018).

39. H. Wickham, ggplot2 (Springer New York, 2009) https://doi.org/10.1007/978-0-387-98141-3 (December 7, 2020).

40. J. Chong, et al., MetaboAnalyst 4.0: Towards more transparent and integrative metabolomics analysis. Nucleic Acids Research 46, W486–W494 (2018).

41. M. Kanehisa, S. Goto, KEGG: Kyoto Encyclopedia of Genes and Genomes. Nucleic Acids Research 28, 27–30 (2000).

42. M. Kutmon, et al., PathVisio 3: An Extendable Pathway Analysis Toolbox. PLOS Computational Biology 11, e1004085 (2015).

43. J. Schindelin, et al., Fiji: An open-source platform for biological-image analysis. Nature Methods 9, 676–682 (2012).

